# *Pseudomonas* can survive bacteriocin-mediated killing via a persistence-like mechanism

**DOI:** 10.1101/719799

**Authors:** PP Kandel, David A. Baltrus, Kevin L. Hockett

## Abstract

Phage tail-like bacteriocins (tailocins) are bacterially-produced protein toxins that can mediate competitive interactions between co-colonizing bacteria. Both theoretical and empirical research has shown there are intransitive interactions between bacteriocin-producing, bacteriocin-sensitive, and bacteriocin-resistant populations, whereby producers outcompete sensitive, sensitive outcompete resistant, and resistant outcompete producers. These so-called ‘rock-paper-scissor’ dynamics explain how all three populations can be maintained in the same environment, without one genotype driving the others extinct. Using *Pseudomonas syringae* as a model system, we demonstrate that otherwise sensitive bacterial cells have the ability to survive bacteriocin exposure through a physiological mechanism. This mechanism is similar to the persister phenotype that allows cells to survive antibiotic exposure, without acquiring antibiotic resistance. We show that a significant fraction of the target cells that survive a lethal dose of tailocin did not exhibit any detectable increase in survival in subsequent exposure (i.e. they survived through a persistence-like mechanism). Tailocin persister cells were more prevelant in stationary rather than log phase cultures. Of the fraction of cells that gained detectable tailocin resistance, there was a range of resistance from complete (insensitive) to incomplete (partially sensitive). By genomic sequencing and genetic engineering we showed that a mutation in a hypothetical gene containing 8-10 transmembrane domains causes tailocin high-persistence and genes of various glycosyl transferases cause incomplete and complete tailocin resistance. Importantly, of the several classes of mutations, only those causing complete tailocin resistance compromised host fitness. This result, combined with previous research, indicates that bacteria likely utilize persistence as a means to survive bacteriocin-mediated killing without suffering the costs associated with resistance. This research provides important insight into how bacteria can escape the trap of fitness trade-offs associated with gaining *de novo* tailocin resistance, and expands our understanding of how sensistive bacterial populations can persist in the presence of lethal competitors.

## Introduction

Diverse microbes inhabit shared environments and compete for limited resources. Competition for these resources can be mediated by secretion of toxins such as antibiotics, type VI effectors, and bacteriocins that enable the producing cells to maintain their dominance (1, 2). Bacteriocins are bacterially produced protein toxins that are lethal toward strains related to the producer (3, 4). These antibacterial toxins have been proposed as antibiotic alternatives to treat or prevent infection spread in both humans and plants (5, 6), in addition to being used in food preservation for several decades (7). Given their ubiquitous nature, where sequenced bacteria are commonly predicted to encode at least one bacteriocin (8–10) and bacteria isolated from diverse environments often produce detectable bacteriocins [eg. (8, 11, 12), it is reasonable to predict that bacteriocin producing populations exert a selective force on co-colonizing sensitive populations to either gain resistance, or avoid killing through a different mechanism. Resistance evolution, often involving a heritable mutation in either the toxin receptor or membrane translocator genes, is a common mechanism to avoid being killed (13, 14). Resistant mutants, however, are likely to suffer fitness costs associated with the mutation, which reduces their ability to proliferate in the environment. Therefore, resistant mutants are only competitive in environments where there is substantial or sustained bacteriocin exposure, otherwise they are competitively inferior to their sensitive progenitors. Conversely, bacteriocin producing populations are dominant over sensitive populations, but are competitively inferior to resistant populations, as the result of the resources wasted on ineffective toxins. These non-transitive dynamics underly a rock-paper-scissor model for microbial competition (13–15), where a community composed of all three genotypes is maintained as a result of negative frequency-dependent selection (14). These dynamics, however, which are built primarily on modelling and laboratory culture-based experiments, are dependent on a small number of qualitative states. It is unknown whether or how quantitative resistance or physiological tolerance in otherwise sensitive populations will affect community competitive dynamics. The ability of bacteriocin sensitive cells to persist under toxin pressure has important implications for the ecology of microbes generally, and for the potential to use bacteriocins as therapeutics specifically. For instance, previous studies have reported that, although sensitive cells may not co-exist with the producing strain in a well-mixed environment (16–18), they still prevail in the competitive *in vivo* environments (13, 14, 19). Little is known, however, regarding the mechanism(s) that allow maintainance of a sensitive population despite sustained bacteriocin exposure.

Bacteriocins are classified into different groups based on their structure, composition, and mode of action. Tailocins are bacteriocins that resemble bacteriophage tails and are grouped into R-type (with a retractile core tube) and F-type (flexible) (20, 21). In the opportunistic human pathogen *P. aeruginosa* and other environmental Pseudomonads and *Burkholderia*, phage-derieved tailocin bacteriocins were shown to antagonize competitors including pathogenic strains (22–28). In fact, a recent study by Principe et al. suggested the effectiveness of foliar sprays of tailocins produced by *P. fluorescens* in reducing the severity and incidence of bacterial-spot disease in tomato caused by *Xanthomonas vesicatoria* (28). Other studies have also indicated the potential use of engineered R-tailocins in suppressing foodborne pathogens using *in vivo* models (29, 30). Our group has previously characterized a R-type tailocin from a plant pathogenic bacterium *P. syringae* pv. *syringae* (*Psy*) B728a (31). This R-type tailocin showed antagonistic potential against several pathovars of *P. syringae* that cause serious diseases and substantial losses in economically important crops such as common bean (pv. *phasiolicola*), soybean (pv. *glycinea*), chestnut (pv. *aesculi*), and kiwifruit (pv. *actinidae*) (31). A broad spectrum of tailocin mediated antagonistic interaction in *P. syringae* has been described recently (11). Yet, we have a very limited understanding regarding the defense responses against tailocin by the target pathogen, a key information to design tailocins as therapeutic agents.

Tailocins are considered to be potent killers as a single tailocin particle is predicted to kill a sensitive cell and an induced producer cell can release as many as 200 particles (32, 33). R-tailocins, once bound to the cell surface receptors of the target cells, puncture through the cell membrane and cause cell death by membrane dissipation (3, 32). Specific lipopolysaccharide (LPS) components of the target cells are known to serve as surface receptors of tailocins (34–36). LPS is composed of three components: the lipid A, core oligosaccharide, and the O-polysaccharide (O-antigen) (37, 38). Although the lipid A and core region are mostly conserved within a species, the O-antigen region varies extensively in its chain length and composition of sugars (39). Biosynthesis and transport of LPS to the outer membrane as well the modification of O-antigen involves complex processes involving a number of highly diverse genes (38). Little is known about the extent of LPS modification required for tailocin resistance and persistence and the consequence of these modifications in host fitness and virulence in the target pathogen.

This study aimed to examine the phenomena that addresses both of the crucial questions related to bacteriocins: 1) how sensitivie cells survive lethal bacteriocin exposure and 2) can bacteriocins be used as stand alone control measures? We demonstrate that a sensitive population can employ multiple strategies to survive toxin exposure without acquiring an otherwise fitness-reducing mutation. One strategy was physiological persistence, a mechanism that enables a sub-population of sensitive cells to transiently survive lethal doses of the bacteriocin without undergoing genetic changes. The second strategy relies on acquiring subtle genetic changes (incomplete resistance) that do not impose a detectable fitness cost, while still allowing the mutants to better survive bacteriocin exposure. We found pronounced differences between the frequencies of persistent cells depending on growth phase of the target cells. Moreover, we identified ten unique mutant alleles with likely roles in the lipopolysaccharide (LPS) O-antigen biosynthesis leading to various degrees of tailocin resistance. We also recovered a mutation in an open reading frame, located in an LPS biosynthesis operon, that results in increased bacteriocin persistence. In addition to increasing the basal persistence frequency, this mutation results in a loss of growth phase-dependent difference in tailocin peristence. Finally, we demonstrated that the complete mutants suffered a fitness cost within a susceptible host plant, whereas both high persisteter, as well as incomplete resistant mutants were equally fit compared to the wild-type.

This work suggests that bacterial cells can employ mechanisms to survive antagonistic toxins but still preserve their host colonization potential. This work has important implications for how bacteria can potentially avoid rock-paper-scissor dynamics widely understood to be important in mediating interactions between toxin producing, sensitive, and resistant genotypes and in using bacteriocins as potential therapeutic agents.

## Results

### A sub-population of *Pph* cells survived tailocin by persistence that increased in the stationary state

The relative activity of the purified tailocin was determined to be 10^3^-10^4^ activity units (AU) and 1.25×10^7^-4.25×10^9^ lethal killing units/ml. The minimum inhibitory concentration was estimated to be 100 AU when exposed to ∼10^6^ viable target cells at their logarithmic growth. No loss of tailocin activity was observed for a period of over six months in the buffer (10 mM Tris PH 7.0, and 10 mM MgSO4) at 4^◦^C.

Purified tailocin was used to test its killing effects on stationary and log phase cultures of the *Pph* target cells in a broth environment. After an hour of 100 AU tailocin treatment, a consistent reduction (3.59 ± 0.12 log) in the viable population occurred for logarithmic cultures, while a significantly lower reduction (1.38 ± 0.14 log) occurred for the stationary cultures. Further analysis showed that, upon treatment of equivalent number of viable cells, stationary cells consistently survived 10 to 100-fold more than the logarithmic cells (**Fig 1** and **Fig S1**).

**Fig 1.**
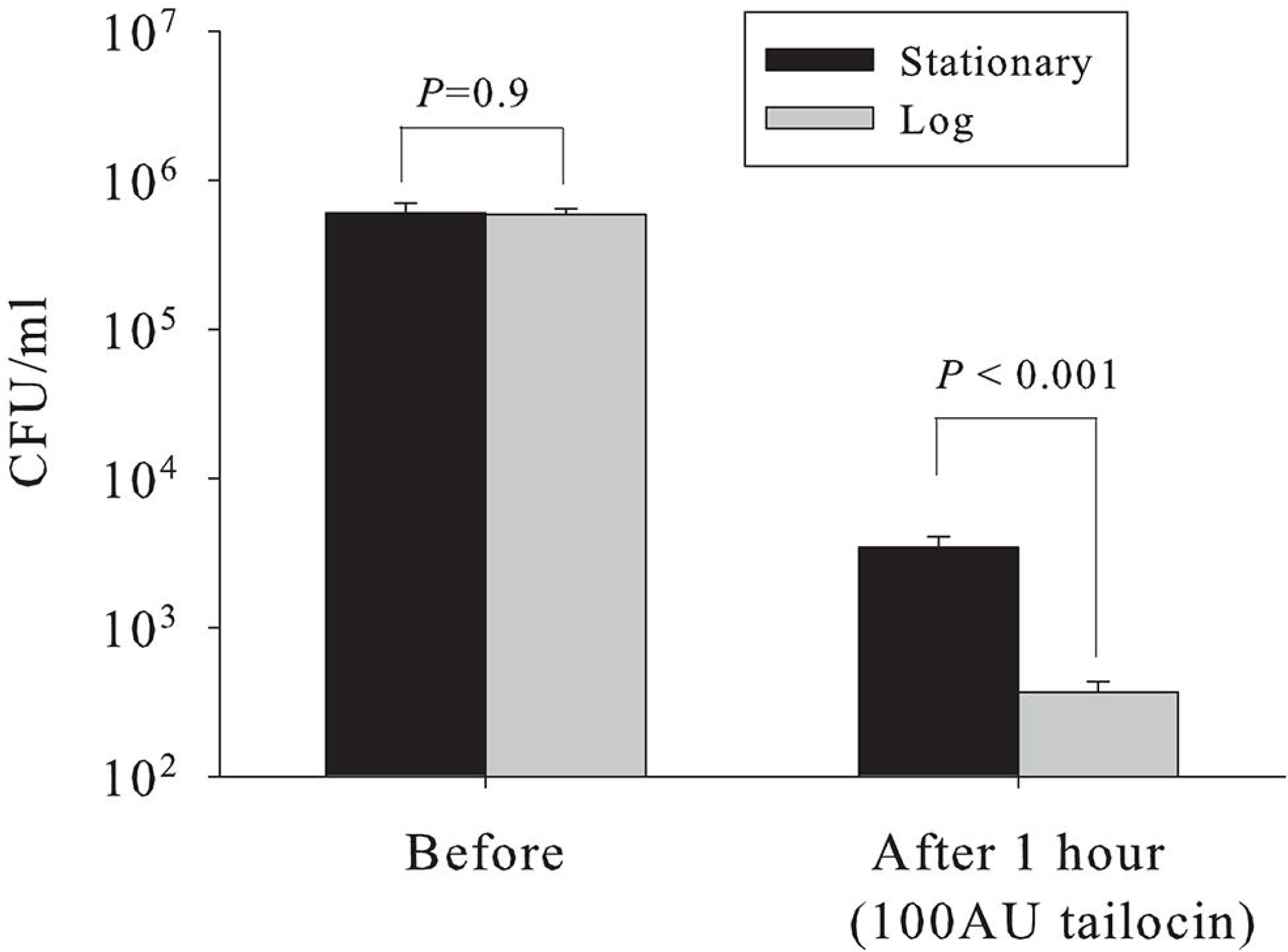
Difference in tailocin survival between stationary and log phase cultures of *Pph*. Cultures were treated with a lethal dose of tailocin (100 AU) and viable cells pre- and post-treatment were enumerated. Experiments were repeated at least five times with 3-6 biological replicates per time. Mean and standard error of mean are graphed. *P*<0.05 indicate significant differences within grouped bars as analyzed in SAS 9.4 with proc Glimmix.

Surviving colonies, especially those from the stationary phase, were predominantly sensitive upon tailocin re-exposure suggesting survival by persistence mechanism (see below).

### The persistent sub-population was maintained under prolonged exposure time and increased concentration of tailocin

Tailocin treatments were applied to both stationary and log cultures of *Pph* for up to 24 hours with enumeration of surviving population before and after 1, 4, 8, and 24 hours of tailocin treatment to generate a tailocin death curve. After a steep reduction in the population within the first hour of treatment, further killing of the cells that survived the first hour treatment, did not occur in either culture (**Fig 2A**). Twenty four hours post-treatment, although the overall population increased (**Fig 2A**), individual treatments showed different results: for some replicate treatments, the population remained constant suggesting maintenance of the persistent state, while for some other replicates, population increased due to replication of cells that acquired tailocin resistance (see **Fig S2**).

**Fig 2.**
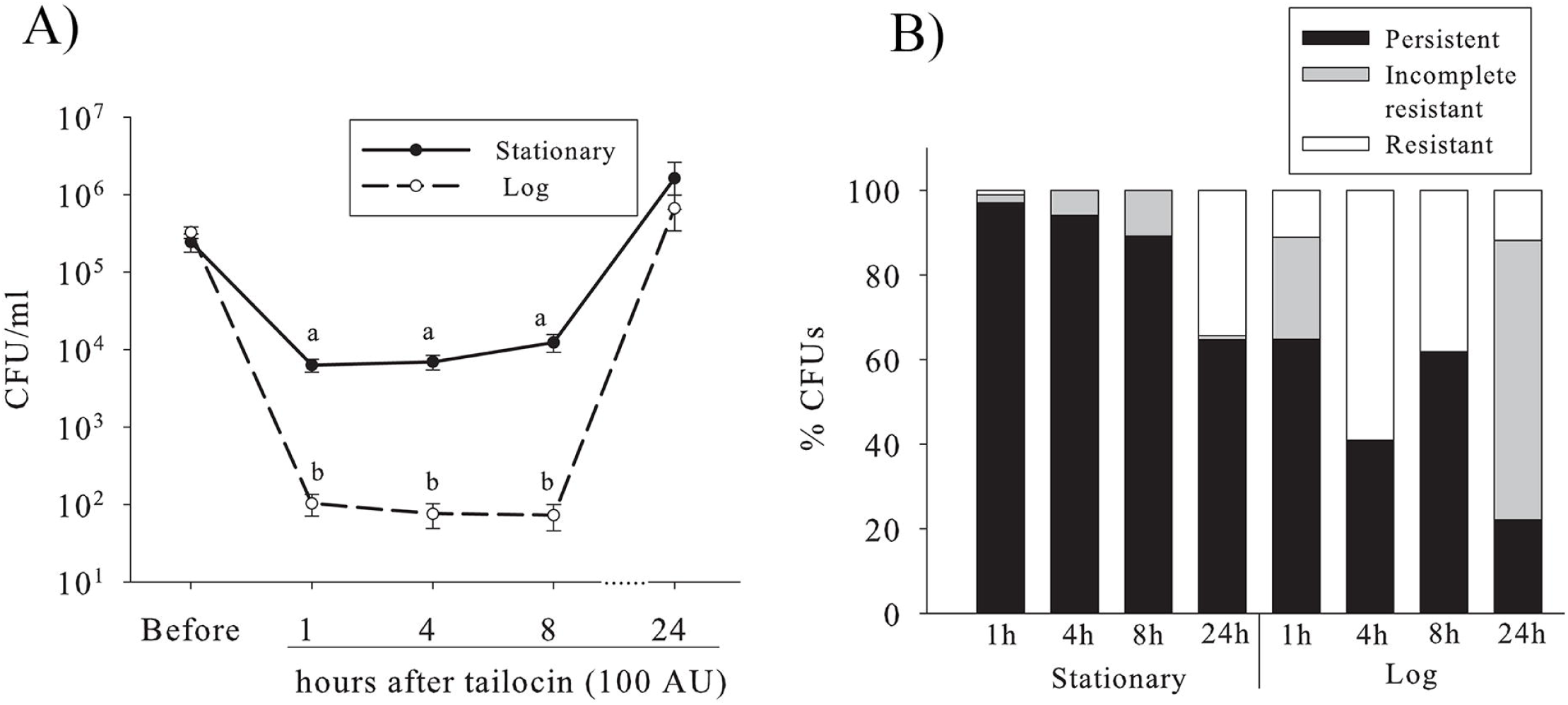
Time dependent dynamics of tailocin survial. **A)** Stationary and log cultures of *Pph* were treated with tailocin (100 AU) and the viable population was enumerated before and 1, 4, 8, and 24 hours post-treatment. Three independent experiments were performed with 3-6 biological replicates per time. Mean and standard error of mean are graphed. Different letters represent significant differences as analyzed in SAS 9.4 with proc Glimmix at *P*=0.05. **B)** Percentage of surviving colony phenotye upon tailocin re-treatment. Randomly selected surviving colonies (n=12-44 for each growth phase and hours of treatment) from the initial tailocin treatment were sub-cultured and re-treated with tailocin, and percentage of the surviving phenotype was calculated. Colonies recovered from three independent experiments were used for the re-treatment to calculate this percentage.

Upon tailocin re-treatment, >90% of stationary and >60% of log cells that survived the first hour treatment, were as sensitive as the wild type (i.e. persistent) as in (**Fig 2B**). The proportion of persistent survivors was higher in the stationary cultures than in the log cultures at all time points (**Fig 2B**). Tailocin persistent cells were recovered from both cultures even after 24 hours of tailocin treatment, although the proportion decreased over time (**Fig 2B**). Tailocin activity was detected in the supernatants recovered from the treated samples that contained persistent cells (**Fig S3**), confirming saturation of tailocin in the treatment. Although a slight reduction of activity was observed when the tailocin preparation was mixed with undiluted stationary supernatant compared to log supertanant, no difference was detected upon diluting the supertants up to 1,000-20,000-fold before mixing with tailocin (similar to how the cultures were diluted for tailocin treatment) (**Fig S4**). This suggested that the increased tailocin persistence in the stationary phase is not related to inhibition of tailocin activity by an extracellular component.

Upon treating the cells with a concentrated tailocin (900 AU) the surviving population decreased such that no difference in survival between the stationary and log cultures was detected (**Fig 3A**). However, even with this higher level of tailocin applied, the proportion of tailocin persistent cells remained higher for stationary phase survivors than that for the log phase survivors (**Fig 3B**).

**Fig 3.**
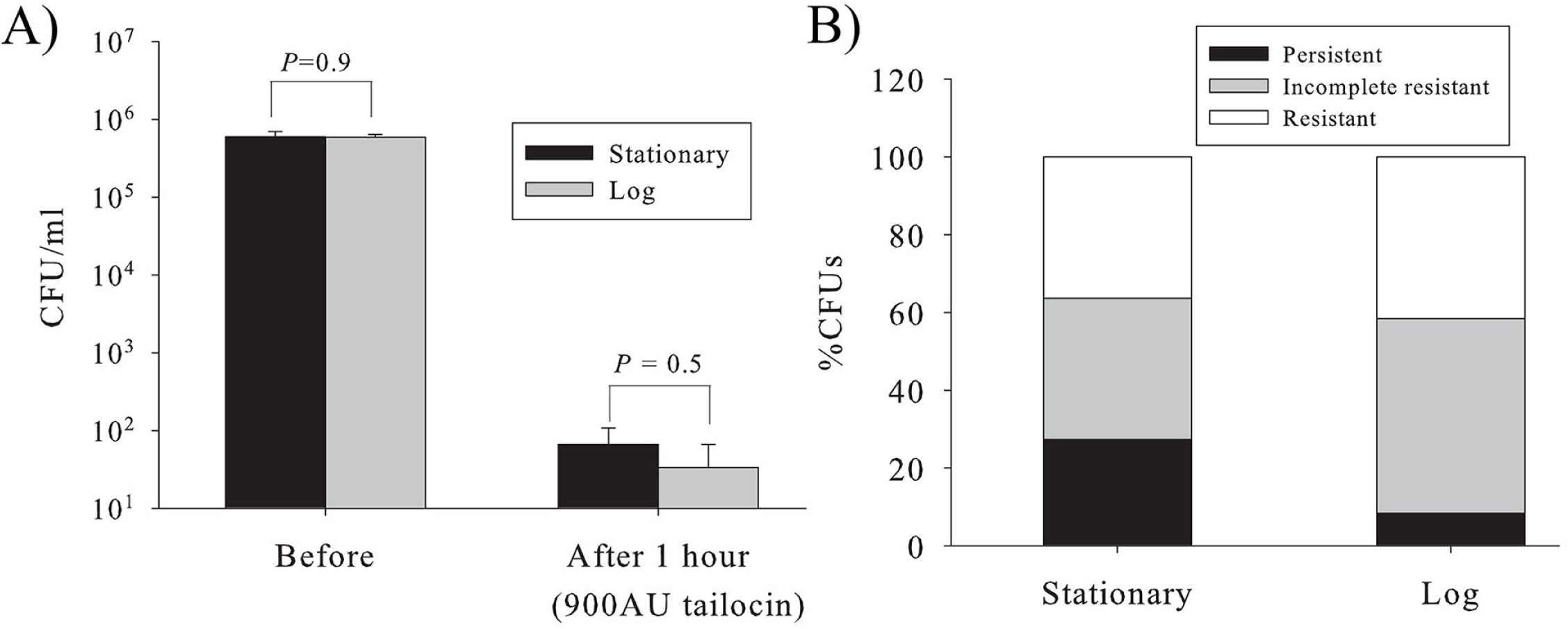
Dynamics of tailocin survival with concentrated tailocin treatment. **A)** Cultures were treated with high dose of tailocin (900 AU) and viable populations pre- and post-treatment were determined. Experiments were repeated at least three times with 3-6 biological replicates per time. Mean and standard error of mean are graphed. *P*<0.05 indicates significant difference within grouped bars as analyzed in SAS 9.4 with proc Glimmix. **B)** Percentage of surviving colony phenotype after treatment with concentrated (900 AU) tailocin for one hour. Although most of the surviving colonies were either incomplete resistant or resistant, persistent cells were maintained even at high tailocin concentration. Surviving colonies were tested during three independently repeated experiments for both cultures. (Refer **Fig. S3** for visual details that leads upto this dynamics.)

### Tailocin exposure selected for heritable mutants showing increased persistence and heterogenous resistance

In addition to the recovery of tailocin persistent sub-population, we recovered an unique mutant, refered here as high persistent-like (HPL), which survived significantly greater than the wild type under liquid-broth treatment (**Fig. 4A**). However, under a long-term exposure of overlay condition, it showed similar sensitivity to the wild type (**Fig. 4B**). Furthermore, the HPL phenotype did not differ in survival between the stationary and log phases even at a higher concentration of tailocin applied (**Fig 5**). Next category of mutants recovered were conditionally-sensitive and are referred here as incomplete resistant (IR) mutants (see **Fig 2B** and 3B for proportion). These mutants lost sensitivity in the broth even at high tailocin concentration (**Fig. 4A**), but displayed some sensitivity in the overlay (**Fig. 4B**). Lastly, complete tailocin resistant (R) mutants that were insensitive to tailocin under both treatment conditions were also recovered. Of the four complete resistant mutants we selected, two (R1 and R4) showed an unique rough colony morphotype.

**Fig 4.**
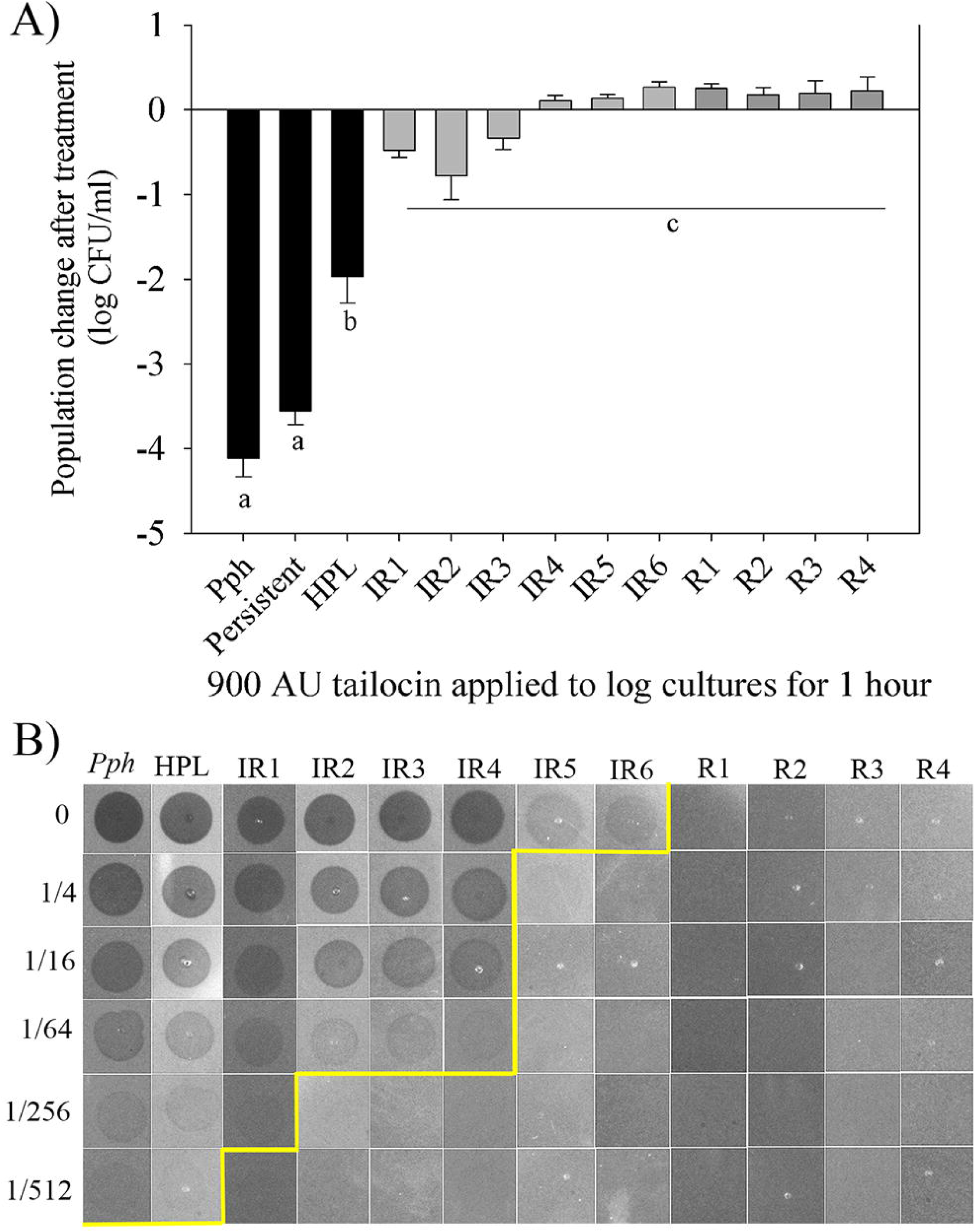
Treatment response of tailocin persistent and resistant lines. A) Reduction in the population of tailocin persistent and resistant mutant lines upon re-treatment with tailocin. Log cultures of each lines were treated with 900 AU of tailocin and change in the population was calculated after an hour of tailocin treatment. At least three separate colonies of each lines were tested and experiments were repeated a minimum of three times. Means of the difference in log transformed viable population pre- and post treatment are graphed. Error bars indicate stantard error of mean. **B)** Assessement of response of mutant lines to tailocin under overlay condition. Dilutions of tailocins (shown on the left most column) were spotted over the culture lawn of each of the lines. Yellow line indicates the dilution upto wchich visible killing was observed. HPL; high persistent-like, IR; incomplet resistant, R; resistant.

**Fig 5.**
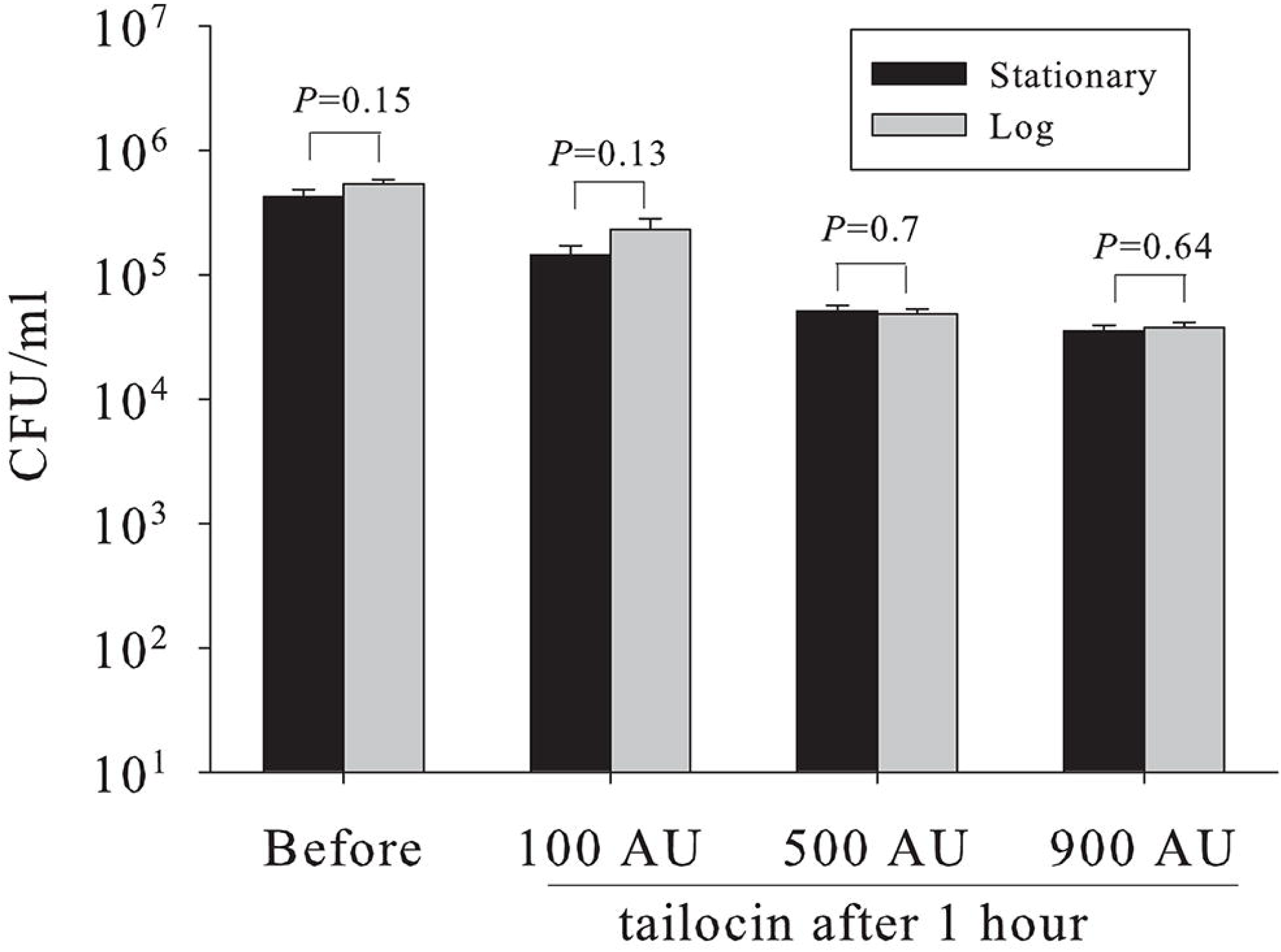
Dynamics of tailocin survival for the high persistent-like (HPL) mutant. Cultures were treated with various tailocin doses and viable cells pre-and post-treatment were enumerated. Means and standard error from three independently repeated experiments with at least three biological replicates per experiment are reported. More killing occurred at high tailocin concentration and no difference in stationary and log phage was observed. *P*>0.05 indicate no significant differences within grouped bars as analyzed in SAS 9.4 with proc Glimmix.

### Mutations involved various genes likely associated to LPS biogenesis and modification

Genome sequencing and variant identification of the high-persistent like (n=1), incomplete resistant (n=9), and resistant mutants (n=4) was performed by mapping the Illumina reads with the parental reference sequence. Mutants isolated at different experiments showed mutation in a different locus. A specific region (**Fig 6**) was identified in the *Pph* genome that showed the most prominent role in tailocin activity. The HPL mutant contained a 16 bp deletion in an ORF that caused frameshift near the C-terminal of a hypothetical protein that is co-transcribed with the LPS genes. No functional evidence could be found for the hypothetical protein by *in silico* analysis except that it was predicted to contain 8-10 transmembrane domains. Majority of genes identified for complete and incomplete resistance were glycosyl transferases and related proteins that are likely involved in LPS biogenesis (**Table 1** and **Fig 6**). For example, for the gene PSPPH_0957 that encodes a glycosyl transferase family 1 protein, a frame-shift and a missense mutation in the middle of the gene caused complete resistance while insertion of few bases at the 3’end of the gene caused incomplete resistance. Moreover, one of the incomplete resistant mutant class (IR6) showed mobilization of the 100 bp MITE sequence present in the *Pph* genome as described by Bardaji et al (40). The insertion of MITE inactivated a gene that is annotated to encode a FAD-dependent oxidoreductase.

**Fig 6.**
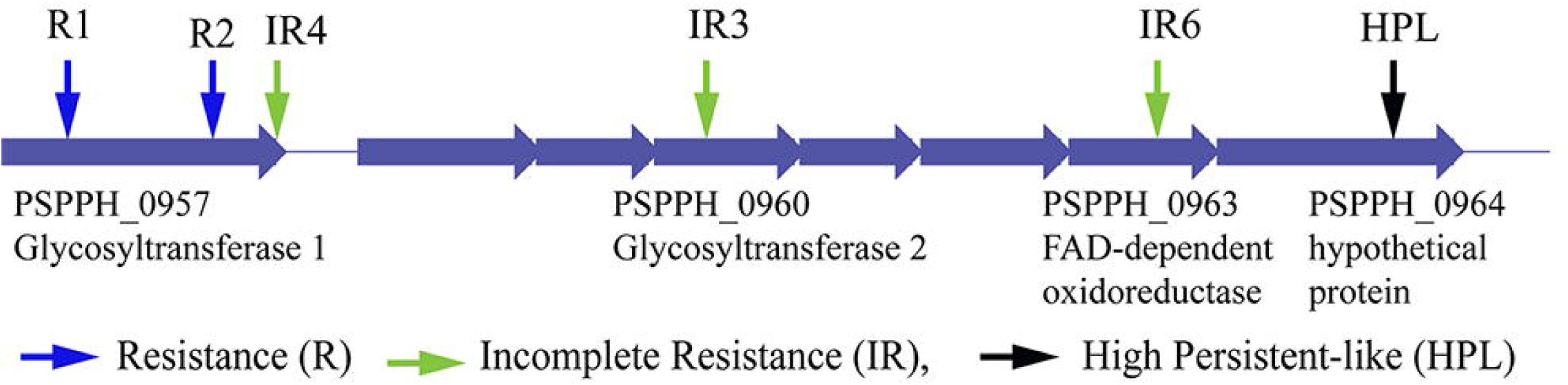
A genomic region (PSPPH_0957-PSPPH_0964) of *Pph* that showed prominent effect on tailocin sensitivity and resistance. Mutations of this region gave rise to multiple phenotypes that differed in tailocin persistence and resistance. Refer Table 2 for a complete list of genes at this and other genomic locations.

**Table 1.**
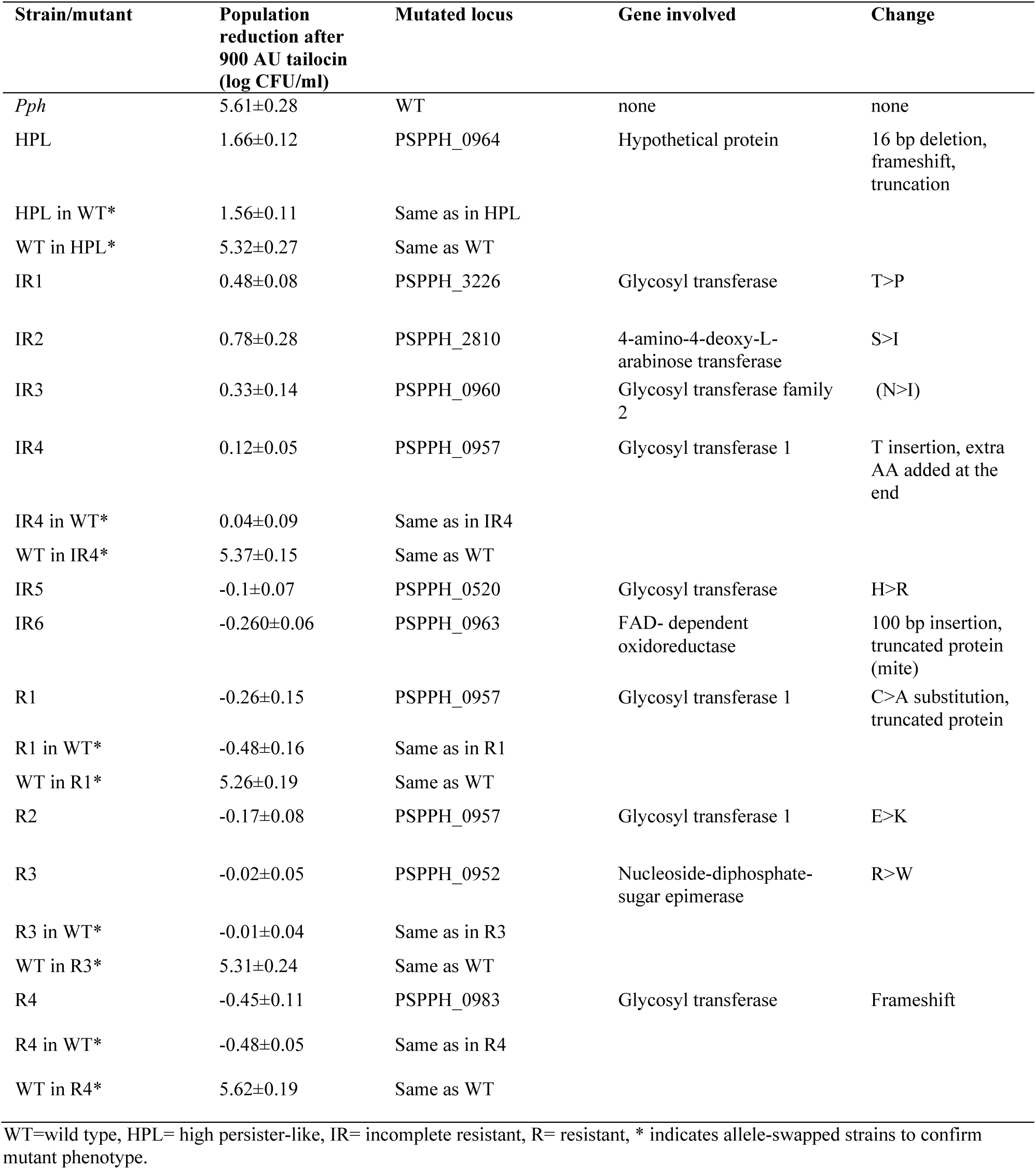
Tailocin sensitivity, predicted loci of mutation and effect on target gene of tailocin high-persistent and resistant mutants and selected allele-swapped strains

**Table 2.** Bacterial strains and plasmids used in this study

| Strain | Background | Characteristics | Source |
| --- | --- | --- | --- |
| B728a | *P. syringae* pv. *syringae* (*Psy*) | Tailocin producing WT strain | (78) |
| B728a:ΔRrbp | *Psy* | A tailocin deficient B728a mutant | (31) |
| 1448a | *P. syringae* pv. *syringae* (*Pph*) | A tailocin sensitive WT strain | (74) |
| ΔhrpL:*Pph* | *Pph* | Type III secretion mutant of *Pph* | (77) |
| S17-1 | *Eschericia coli* | Strain that can conjugate *P. syringae* |  |
| pDONR207 | plasmid | Gate way entry clone (Gm<sup>R</sup>) | Invitrogen |
| pMTN1907 | plasmid | Gateway compatible destination vector (Km<sup>R</sup>,Tetr, Sucs) | (77) |
| pDONR1K18ms | plasmid | Gateway vector (Km<sup>R</sup>,Cm<sup>R</sup>, Sucs) | Unpublished<br>Gifted by Dr. Brian Kvitko, University of Georgia |

The mutant phenotypes detected by genome sequencing were further confirmed by sanger sequencing of the target region as well as by swapping the mutant alleles to the wild type background and vice-versa of the selected mutants (HPL, IR4, R1, R3, and R4). In all cases, the allele-swapped strains showed the phenotypes as expected (**Table 1**). This confirmed that the mutation identified by genome sequencing were responsible for the tailocin high persistent-like and resistant phenotypes.

LPS analysis of the mutants and the wild type *Pph* showed that the complete resistant mutants lacked fully-formed O-antigen region, whereas the high persistent-like and majority of incomplete resistant mutants still possessed the O-antigen with minor to undetactable changes. One of the incomplete resistant mutants (IR4), however, showed a very different and faint O-antigen band (**Fig S5**).

### *In planta* fitness was compromised only for complete resistant mutants devoid of LPS O-antigen

As indicated by the population levels of each mutant lines in the green bean plants, while the high persistent-like and incomplete resistant mutants did not suffer any, the complete resistant mutants suffered a significant fitness cost as earley as 24 hours post-inoculation (**Fig. 7** and **Fig. S6**). Up to a 100-fold reduction in the population was seen for the complete resistant mutants (equivalent to the reduction in the type III secretion system mutant). At 48 hpi, although the complete resistant mutants population increased compared to the type III mutant, it was still significantly lower than wild type, HPL and IR mutants (**Fig. 7** and **Fig. S6**). These results suggest that persistence and incomplete resistance are mechanisms to survive attack by the competitor strains while keeping the host fitness and virulence potential intact.

**Fig. 7.**
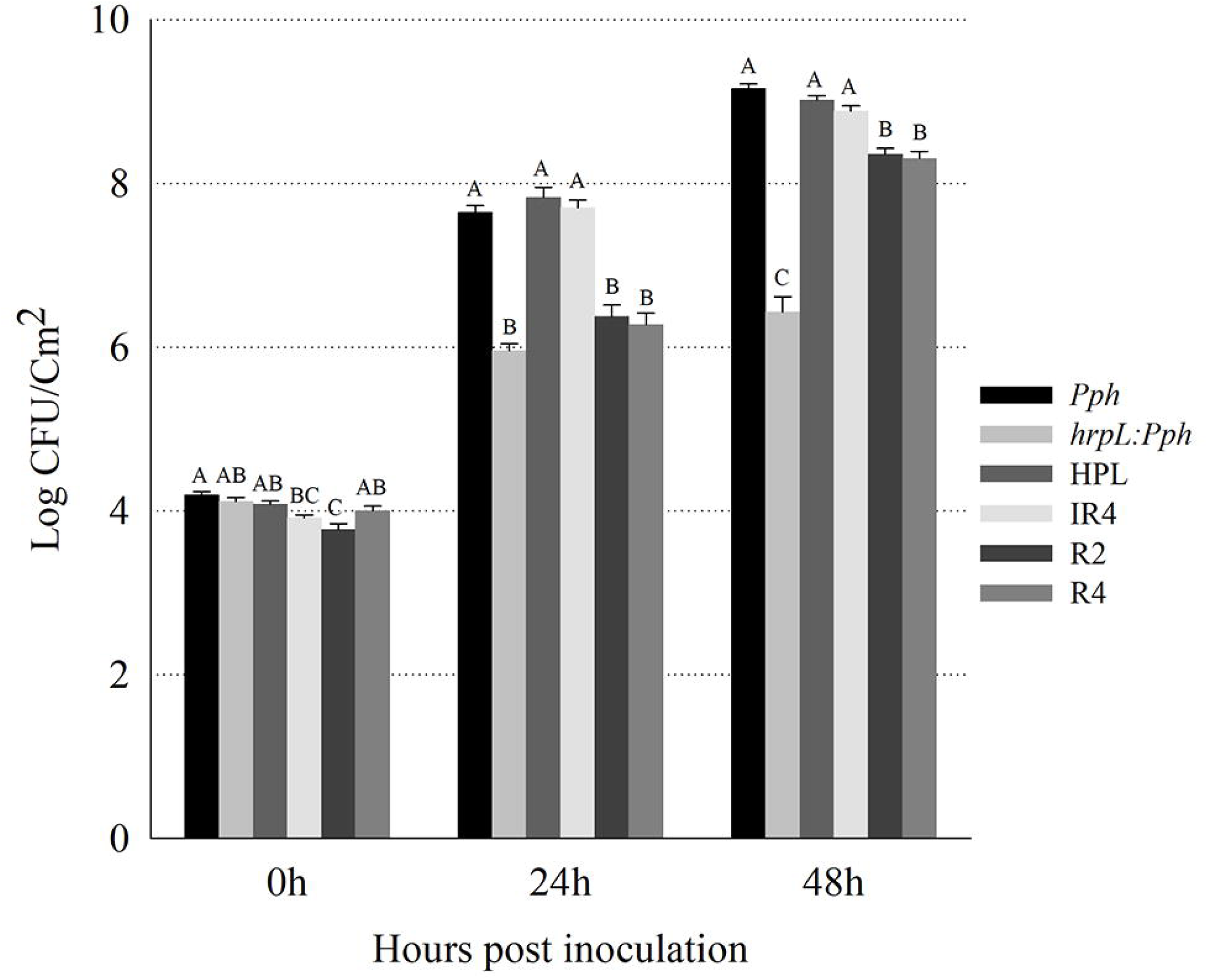
*In planta* characterization of tailocin high persistent, selected incomplete resistant, and resistant mutants. Strains and mutants were syringe infiltrated into green bean leaves and population change was monitered by harvesting infiltrated leaf discs and enumeration using dilution plating. Experiments were repeated atleast twice with 8 replications per strain. A representative experiment is presented. Error bars indicate standard error. Different letters indicate significant difference (*P<*0.05) for a given time point.

## Discussion

There have been renewed research interests in alternative treatment strategies for bacterial pathogens due mainly to the growing threats of antibiotic resistant infections (5, 7). Bacteriocins including tailocins have long been proposed as effective and more specific alternatives to broad spectrum antibiotics (5, 6). However, a critical question that remains unaddressed in designing bacteriocins as pathogen control agensts is; how does the sensitive pathogen population respond to application of lethal doses of bacteriocins? Moreover, although it is predicted in the rock-paper-scissors model of bacteriocin mediated interaction, that sensitive cells co-exist with the producer through ecological mechanisms (13, 14, 19), there is no description of the possibility of maintaining a bacteriocin sensitive population through a physiological mechanism.

In this study, we addressed these questions using a phage-tail like bacteriocin (i.e. tailocin) produced by *P. syringae* pv. *syringae* strain B728a in killing target cells of *P. syringae* pv. *phaseolicola* 1448A. We showed that, upon exposure of a lethal dose of tailocin, a sub-population of sensitive cells survives without undergoing genetic changes. The fraction of this sub-population, termed here as tailocin persistent sub-population, increased significantly in the stationary phase than in the logarithmic phase of growth. By repeated exposures of this sub-population to same or higher doses of tailocin, we showed that they have not gained any heritable resistance, and physiological persistence is the only mechanism for their survival.

Moreover, a prolonged tailocin exposure generated a killing pattern similar to that reported for persistent sub-population upon antibiotic treatment (41, 42). Persistence was maintained for at least 24 hours with tailocin exposure, a phenomenon that was more evident in some treatment replicates in which resistant evolution did not occur (see Fig. S2). Although increasing the tailocin concentration killed some of the persistent survivors, and the difference in survival between the two growth phases was no longer seen, stationary phase-derived cells still exhibited higher persistence than the log phase cells upon re-exposure. This indicated that the stationary cells may require multiple hits by tailocin particles as opposed to the one-hit-one-kill mechanism of killing described for tailocins (32, 33), or that the probabilty of a successful hit in stationary phase is lower than in log phase. Since tailocins are thought to be target cell specific, and are not known to have off-target effects, higher concentration of tailocin could be used to achieve a more effective pathogen control. However, although at a low level, persistence was still maintained even with high-dose tailocin treatment and inherent emergence of either complete or incomplete resistance was frequently observed. As such, although a significant reduction in pathogen population and disease pressure can be obtained with tailocins, a stand-alone tailocin treatment might not be enough to achieve a sustainable pathogen control.

The use of the term ‘persistence’ in relation to antimicrobial survival is disputed to some extent and is sometimes used interchangably with ‘tolerence’ and ‘viable but not culturable state’. In this paper, we used ‘persistence’ as this phenotype was only seen in a sub-population, resulted in a bi-phasic death curve, cells resuscitated almost immediately upon tailocin removal, and were equally sensitive to the wild type cells upon re-exposure. This definition of persistence has been suggested previously (42). Persistence to antimicrobials is being increasingly recognized for its role in antimicrobial treatment failures with bacterial infections (43). Various mechanisms are implicated in the maintainance of persistence (44, 45), although persistence responses can be different based on the stresses involved and their mode of action (46). Of these mechanisms, toxin-antitoxin (TA) systems are the most studied ones for formation of persister-subpopulation (47–49). TA systems were shown to be induced when cells were starved for certain sugars and amino acids or by exposure to osmotic stresses that altered ATP levels in the cell (46, 50).

However, It was also shown that activation of TA system does not always induce persister formation (50). Additionally, recent findings have indicated a mechanism mediated by the guanosine penta- or -tetraphosphate (ppGpp) for persister formation that is not dependent on a TA system (51). A strong stationary state effect, that likely involved starvation response, was shown to increase persistence by 100-1,000 fold in *Staphylococcus aureus* with ciprofloxacin treatment (50). Whether similar mechanisms of TA and independent ppGpp systems regulate tailocin persistence or a specific mechanism for tailocin and/or related bacteriophage exists, remains to be determined. Nevertheless, our data of the difference in tailocin persistence between the stationary and log cultures suggests that metabolic inactivity and starvation-induced stress could be a strong factor in tailocin persistence.

Few previous studies have demonstrated growth-phase dependent differences in LPS O-antigen chain length and composition or their regulatory pathways (52, 53). In a previous study with *P. fluorescens,* exponentially growing cells had a significantly higher rate of cell lysis than stationary or decline phase cells with bacteriophage PhiS1 (54). Moreover, a recent study showed that sensitivity to colicin, a bacteriocin from *Eschericia coli*, could be altered by growth condition dependent LPS O-antigen changes (55). Growth phase dependent LPS modification, although has not been reported so far in *P*. *syringae*, could be contributing to the difference in tailocin persistence. We also observed that undiluted stationary phase supernatant inhibited tailocin activity to some extent compared to the log phase supernatant. However, whether this is linked to differences in the secreted LPS between the two supernatants or other cellular factors needs further assessment. Moreover, we also demonstrated that mutation in one of the hypothetical proteins containing a signal peptide and several trans-membrane domains caused increased tailocin persistence. Since the hypothetical protein occurs in the same operon as other LPS biogenesis genes, it is likely that it plays a role in O-antigen biogenesis and/or modification, thereby reducing tailocin interaction with the cells.

Phase variation is another mechanism that is known to cause increased survival to surface active antimicrobials (eg. phages and host immune defenses) (56). Phase variation is a gene regulation system that induces heterogenous expression of specific genes in a clonal population (56–58). Although phase variation is heritable, the ‘ON’ ‘OFF’ switch from variant to wild type phenotype occurs randomly amounting to 10^-4^ to 10^-1^ per generation, significantly greater than what is expected by mutational events (59). Phase variation has been shown to modify LPS operons in *S. enterica* spp (60), *P. aeruginosa* (61), and *P. fluorescence* (62). In *S. enterica* serovar Typhimurium phase variable glucosylation of the O-antigen was reported to cause temporal development of phage resistance (63). Other reports of the role of phase variation in defense against bacteriophages, that also target LPS, are available (64, 65), including in the closely related *P. aeruginosa* (61). Therefore, phase variation in tailocin survival can not be ruled out since LPS is the receptor involved. Since the rate of switch from the tailocin surviving variant to the sensitive wild type upon tailocin removal will have to be highly greater than reported (59), the tailocin persistent phenotype is unlikely to be the result of phase variation. It remains to be determined, however, if the incomplete resistance could be caused by phase variable gene switching. In one of the incomplete resistant mutants, the target gene was modified by movement and integration of an internal mobile genetic element (MGE). MGEs have been shown to cause phase variation in other bacterial systems (66, 67). A high-throughput sensitivity screening of the mutant progeny colonies will be required to confirm if the tailocin resistant phenomena involves phase variable gene switch.

To our knowledge, the persistent as well as incomplete resistant phenotypes observed here have not been described before with bacteriocins. Here, using a *Pseudomonas syringae* tailocin, we showed that, in addition to the persistent sub-population, resistant lines with various degrees of sensitivity are selceted by exposure of target cells to tailocins. LPS analysis and *in planta* fitness assessement showed that, to gain complete tailocin resistance, cells have to loose their LPS O-antigen. This comes with a fitness trade-off. On the contrary, by maintaining a persistent sub-population and/or by undergoing subtle genetic changes, the LPS integrity was preserved for the most part and fitness within the host was maintained. Similar results of reduced virulence and plant fitness was shown in phage resistant mutants devoid of O-antigen (68). Moreover, two of the complete resistant mutants (R1 and R4) in our study showed a typical rough-colony morphotype. Rough-colony morphotypes lacking LPS O-antigen were resistant to phages and had reduced *in planta* virulence as described previously in *P. syringae* pv. *morsprunorum* isolates from plum and cherry (62). In other plant pathogenic bacteria such *as Xanthomonas oryzae* pv. *oryzae* and *Xylella fastidiosa,* loss of O-antigen, respectively, reduced type III secretion into plants (69), and increased recognition of the pathogen by the host immune response (70). In both cases, plant virulence of the mutants was significantly reduced (69, 70). Also, the incomplete resistant mutants, although contained mutations in the LPS biogenesis genes and mostly had their O-antigen region intact, did not show any fitness trade-offs in our growth chamber experiments. However, under a natural environment that involves more severe environmental stresses, it could be expected that their fitness would be altered. Under these circumstances, the persister sub-population, that does not involve any genetic changes, would enable the target pathogen to withstand the host defence while surviving the competition and ensure that its lineage is maintained.

Taken together, both previous reports and our results suggest that full resistance to tailocin incurs significant fitness cost. Our work demonstrates that persister and incomplete resistant sub-populations of the sensitive strain preserve their host fitness despite suffering bacteriocin exposure. These results have important implications for the mechanisms and ecological processes that can promote the co-existence of both sensitive and bacteriocin producing populations. In **Fig 8**, we present a model of the three cell types (resistant, incomplete resistant, and persister) that can be detected following bacteriocin-mediated selection. In particular, we view the persister sub-population as one that can survive bacteriocin exposure, without paying any long term fitness cost. In the short term, however, we believe that persister cells have a fitness of zero because they are likely not replicating (see Figure S2). Thus, bacteriocin persisters may best be thought of as a means to switch at high frequency between being phenotypically resistant and phenotypically sensitive.

**Fig. 8.**
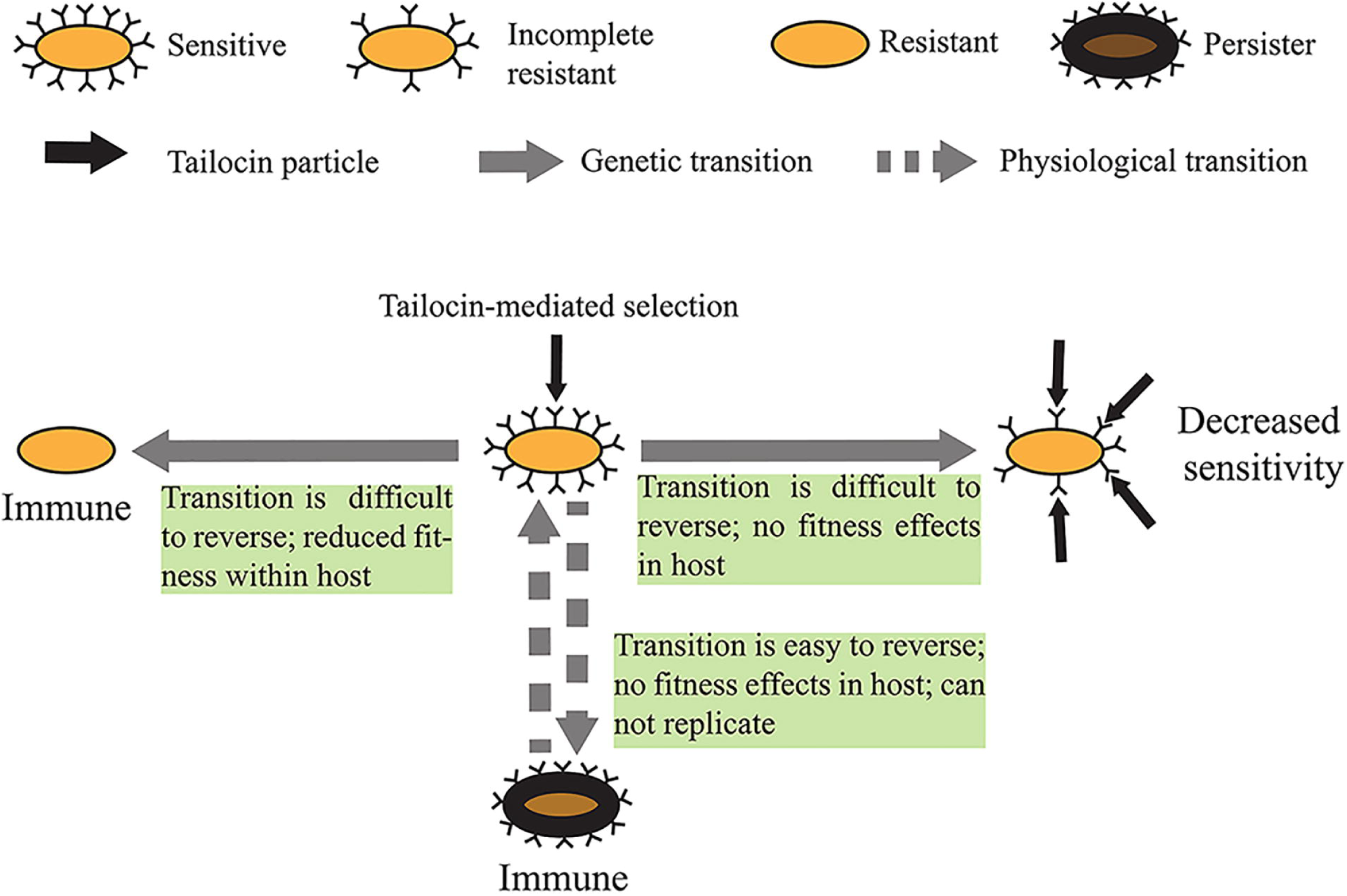
A visual representation of the various tailocin surviving sub-populations and their phenotypes upon exposure to tailocin.

## Materials and Method

### Bacterial strains, media, and culture conditions

All bacterial strains, plasmids, and mutants used in this study are listed in Table 1 and 2. *P. syringae* pv. *syringae* (*Psy*) wild type (WT) strain B728a and its tailocin defective mutant ΔRrbp were used to prepare the treatment supernatants. *P. syringae* pv. *phaseolicola* (*Pph*) 1448A was used as the target strain. Tailocin high-persistent like, resistant, and selected complementated strains of *Pph* generated in this study are described in Table 1. King’s medium B (KB) was used to culture the strains. Liquid cultures were prepared by inoculating individual colonies from a two-day old KB agar plate into 2 ml of liquid medium at 28^◦^C with shaking at 200 rpm.

Antibiotics Kanamacyin (Km), Chloramphenicol (Cm), Tetracycline (Tet), Gentamycin (Gm), Rifampcin (Rif), Nitrofurantoin (NFT), and Nalidixic Acid (NA) were used at 50, 25, 10, 50, 50, 50, 50 µg/ml final concentration, respectively.

### Tailocin induction and purification

Purified tailocin and control treatments were prepared from logarithmic (log) cultures of *Psy* B728a and ΔRrbp, respectively using a polyethylene glycol (PEG) precipitation protocol as previously described (31, 71). Briefly, 100-fold diluted overnight B728a cultures were sub-cultured for 4-5 hours in KB before inducing with 0.5 µg ml^-1^ final concentration of mitomycin C

(GoldBio). Induced cultures were incubated for 24 hours with shaking at 28^◦^C. Next, cells were pelleted by centrifugation and the supernatants were mixed with 10% (w/v) PEG 8000 (FisherScientific) and 1M NaCl. Supernatants were then incubated either in ice for 1 hour and centrifuged at 16,000 g for 30 min at 4^◦^C or incubated overnight at 4^◦^C and centrifuged at 7000g for one hour at 4^◦^C. Pellets were resuspended (1/10-1/20^th^ of the original volume of the supernatant) in a buffer (10 mM Tris PH 7.0, and 10 mM MgSO4). Two extractions with equal volume of chloroform were performed to remove residual PEG. Tailocin activity was confirmed by spotting dilutions of 3-5 µl of both the tailocin and control supernatants onto soft agar overlays of *Pph*. The relative activity of tailocin was expressed as arbitrary units (AU) as obtained from the reciprocal of the highest dilution that exhibited visible tailocin killing in an overlay seeded with ∼10^8^ CFUs of *Pph* log cells. Lethal killing units of the purified tailocin were determined using a Poisson distribution of the number of surviving colonies after treatment with different dilutions of the tailocin as described previously (27, 72).

### Tailocin treatment and survival assessment for stationary and log cultures

To assess tailocin activity against the stationary and log phases of *Pph*, individual colonies growing on KB agar plates for ∼2 days were inoculated into 2 ml of liquid KB medium. Following incubation at 28^◦^C with shaking at 200 rpm overnight, the cultures were back diluted 1000-fold into fresh KB. The back diluted cultures were either incubated for 28-30 hours to prepare stationary cultures, or back-diluted 100-fold at 24 hours and cultured for another 4-6 hours to prepare log cultures (**see Fig S7** for a growth curve of *Pph*).

Stationary cultures were diluted 20,000^-^fold and logarithmic cultures were diluted 1,000-fold [∼10^5^ −10^6^ CFUs/ml for both cultures see **Fig 1**] in fresh KB before tailocin treatment. Treatment was applied by mixing 10 µl of diluted cultures in 90 µl of purified tailocin diluted in KB. After treatment, samples were incubated for ∼ 1 hour at 28^◦^C and washed twice to remove residual tailocin particles. Washing was performed by mixing the treated culture in 900 µl of fresh KB followed by centrifugation at 12,000 g for 2 min. The top 900 µl fraction was discarded and the bottom 100 µl fraction was serially diluted and either spread- or spot-plated to enumerate surviving population. Plates were incubated at 28^◦^C for 1-3 days before enumeration. Serial dilutions of both stationary and log cultures were spotted onto KB agar to enumerate the untreated population. Experiments were performed with various tailocin concentrations (i.e. 100 AU, 500 AU, and 900 AU).

### Tailocin re-treatment to differentiate persistence and resistance

Surviving colonies were treated again with tailocin to differentiate them into persistent or resistant colonies. Re-treatment was performed by an overlay method as described previously (71), or by broth treatment as discussed above. Overlay method was used to determine the AU of the tailocin preparation with the selected mutant lines. Broth exposure was used to calculate reduction in the population of log cultures after treatment. Surviving colonies were differentiated into various phenotypes as follows: persistent (sensitive to tailocin to the wild-type level in both the overlay and broth method), high-persistent like (completely sensitive in the overlay but survived significantly more than wild-type under broth condition), incomplete resistant [showed conditional sensitivity (i.e. some sensitivity in overlay but no significant sensitivity in the broth), and complete resistant (were insensitive under both conditions).

### Time dependent death curve with tailocin treatment

Prolonged tailocin exposure was performed with both the stationary and log-phase cultures of *Pph* to generate death curves. Treatment was applied in a 96-well plate as before. Surviving populations were enumerated at 1, 4, 8, and 24 hours following treatment and randomly selected surviving colonies were re-exposed to tailocin to differentiate them into persistent or resistant phenotypes.

### Tailocin recovery from the treated samples and activity testing

Stationary and log cultures treated with purified tailocin for 1-24 hours as described above were centrifuged and the supernatant was collected, and filter sterilized using a 0.22 µm syringe filter. Supernatants were diluted 5-, 10-, 50-, and 100-fold in KB and spotted on *Pph* overlay. Purified tailocin particles diluted in KB was also included as control treatment.

### Determining the effect of stationary and log supernatant on tailocin activity

Stationary and log phase cultures were prepared as described above by culturing *Pph* cells in KB broth for either 28-30 hours or for 4-6 hours, respectively. Cultures were centrifuged for 2 min at 12,000 g and the supernatant was filter sterilized using a 0.2 µm syringe filter. Stationary and log supernatants were diluted 1,000-20,000-fold in KB (according to how the cultures were diluted for tailocin treatment). Various dilutions (10-, 50-, 100-, and 1000-folds) of purified tailocin were prepared in the stationary and log supernatants. Dilutions were spotted on a *Pph* lawn using the overlay method.

### Genome sequencing and analysis of the tailocin high-persistent and resistant mutants

The tailocin high-persistent like (HPL) and complete and incomplete resistant mutants recovered from tailocin treatment of *Pph* wild type cells (and confirmed by re-treatment) were selected for genome sequencing. DNA was extracted from the overnight cultures using Promega Wizard Genomic DNA purification Kit using manufacturer’s protocol. DNA quantity and quality were assessed with Qubit 3 Fluorometer using Qubit™ dsDNA HS Assay Kit (Thermo Fisher Scientific) and Nanodrop 2000 (Thermo Scientific). A uniquely indexed library from each mutant line was prepared using Illumina DNA Flex kit (Illumina). An approximately equimolar pool of libraries was generated and 150bp paired-end reads were sequenced on an Illumina MiSeq.

Resulting forward and reverse read files were paired and mapped to the *Pph* 1448A chromosomal and plasmid sequences using Geneious R10.2 with default parameters for medium sensitivity. Next, genetic variants were identified using ‘Find variations/SNPs’ program within Geneious using default settings. Regions supported by a minimum of 10 reads and >90% variant frequency were selected. Moreover, variants shared among all the mutants that were generated in independent experiments were discarded as misalignments. Variants identified by Geneious were confirmed in the contigs generated by De Novo assembly of the paired reads using SPAdes 3.11.0 using K-mer sizes of 21, 33, 55, 77, 99, 127 with careful mode selected to minimize mismatches and short indels. Indels were also detected and visualized in the contigs by Harvest suit tools (73). Sequencing generated 1,986,400 −2,857,324 paired reads per genome with 48-69X genome coverage. *De novo* assembly generated 308-338 contigs per genome with the N50s of 75,222-81,220bp. The total assembly size was 5.97 Mbp with a GC content of 57.96%, values similar to *Pph* reference genome (74). The read files of all the genomes generated in this study have been deposited to the NCBI Short Read Archive (SRA) database under a Bioproject PRJNA608702 and biosamples SAMN14206700-SAMN14206710. The SRA accession numbers are SRR11179092-SRR11179102.

Presence absence and similarity searches of the selected genomic regions and genes implicated in tailocin persistence and resistance were performed with NCBI and IMG-JGI databases using BLAST algorithm using the *Pph* sequences as query. InterProScan (75) and Phobious program within the Geneious plugin was used to predict functional domains in the amino acid sequences.

### Confirmation of mutant phenotype by allele swap

The mutations were further confirmed with sanger sequencing and by swapping the mutant allele to the wild type background and vice-versa. Allele swap experiments of the selected incomplete resistant (IR4) and complete resistant mutants (R1, R3, and R4) were performed as previously described in Hockett et al (31). Briefly, a ∼1kb fragment containing the mutant or a WT allele was amplified using a Phusion High-Fidelity DNA Polymerase (New England Biolabs) using standard protocols. Primers used for generating the PCR fragments and mutant confirmation are listed in Table 3. The PCR fragment contained gateway cloning sites added through the primer extension. Purified PCR fragment was cloned into pDONR207 and further recombined into pMTN1907 using BP and LR clonase enzymes, respectively. pMTN1907 containing a desired clone was transformed to S17-1 and conjugated to the *Pph* wild type or mutant background by bi-parental mating. Tet_R_ merodiploids of *Pph* selected on Tet, Rif, and NFT plates were counter-selected in KB supplemented with 10% Sucrose, followed by PCR confirmation of the desired allele-swapped strains. Allele swap of the high persistent-like (HPL) mutant was performed similarly, but using a one-step gateway vector pDONR1K18ms and the merodiploids were selected in Km, Rif, NFT plates.

**Table 3.**
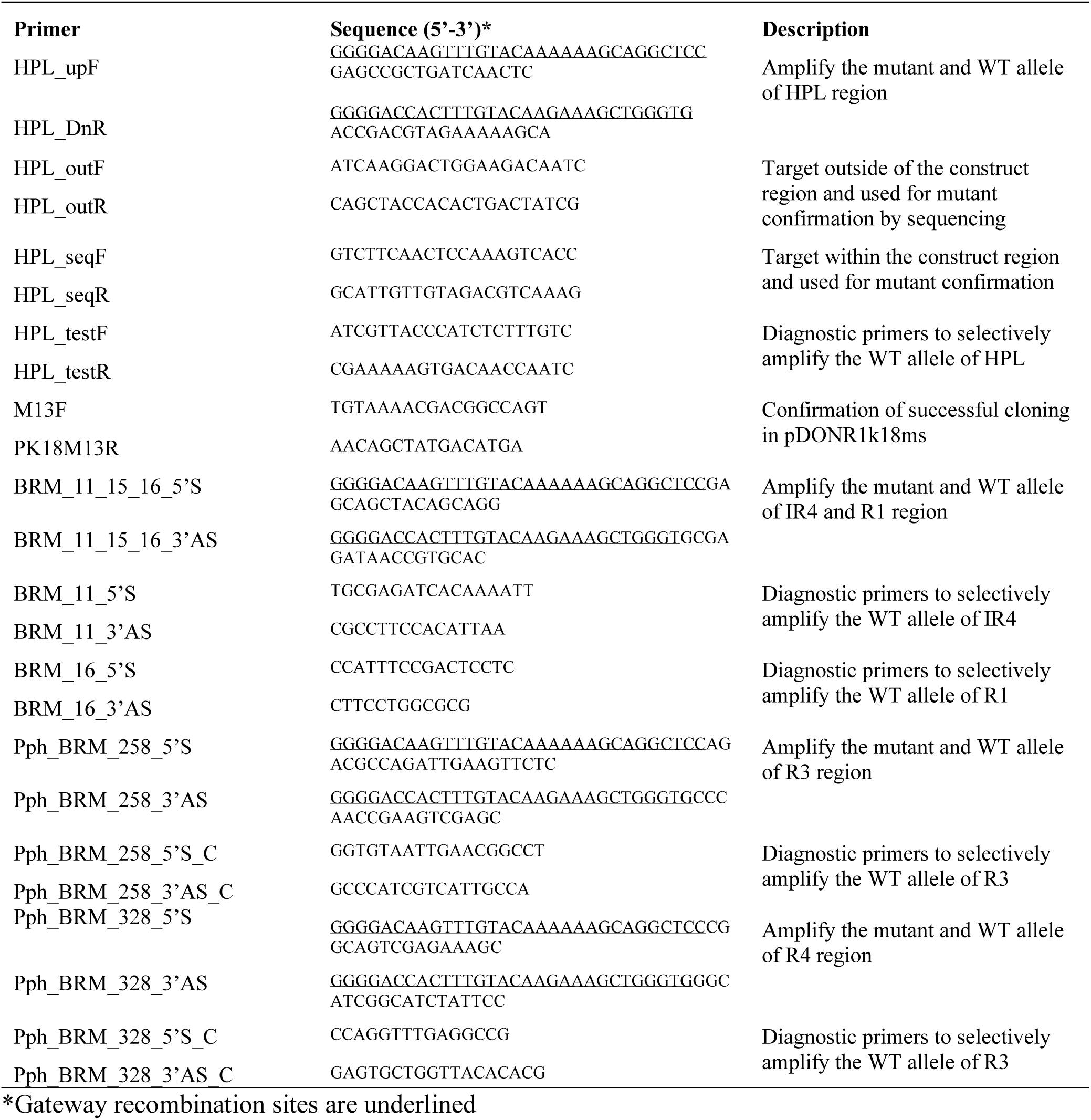
Primers used in this study

### LPS extraction and visualization

LPS extraction was performed as described by Davis and Goldberg (76) from pellets of one milliliter of overnight cultures (OD_600_ 0.5). After extraction, 10 µl samples were separated by SDS-PAGE. Samples were labeled using the Pro-Q Emerald 300 glycoprotein stain kit (ThermoFisher Scientific, #P21857) according to manufacturer’s instructions. For LPS size determination, CandyCane™ glycoprotein molecular weight standards (ThermoFisher Scientific, #C21852) were included. Gels were visualized using Molecular Imager Gel-Doc XR+ (Bio-Rad) with Image Lab Software.

### In planta fitness test of the high-persistent, incomplete resistant, and resistant mutants

The high persistent-like (HPL), selected incomplete resistant, and complete resistant mutant of *Pph* including a type III secrection mutant Δ*hrpL::Pph* (77) were tested for their *in planta* fitness. *In planta* experiment was performed in a growth chamber (Conviron) maintained at 24◦C, 75% RH and 16 and 8 hours of day/night cycles. Plants of Dwarf French Bean (*Phaseolus vulgaris*) variety ‘Canadian Wonder’ were grown in a Dillen 6.0 Standard pots (Onliant) in Sunshine Mix 4 Aggregate Plus Professional Growing Mix (Growerhouse). Plants were irrigated daily. Nine days post seeding, plants were infiltrated with suspension of the bacteria. For inoculant preparation, overnight cultures of WT *Pph*, Δ*hrpL::Pph*, and the tailocin persistent and resistant lines were pelleted by centrifugation, washed twice, and resuspended in equal volume of 10 mM MgCl2 buffer. Optical density (OD_600_) was adjusted to 0.1 using a Spectronic 200 Spectrophometer (Thermo Scientific) and diluted 50 times. ∼200 µl diluted cultures were infiltrated onto designated areas of the two primary leaves using 1 ml BD syringes and infiltrated areas were marked. Infiltrated areas were harvested using a 1cm cork borer in a 2 ml tube containing 200 µl 10 mM MgCl2 and 2 of the 3 mm glass beads (VWR) at 0, 24, and 48 hours post infiltration. Harvested leaf discs were homogenized in a FastPrep-24 instrument (MP Biomedicals) for 20 sec. Homogenate (5 µl) after serial dilution were spotted on KB agar plates supplemented with 50 µg/ml of Nalidixic Acid and CFUs were counted after two days. *In planta* experiments were repeated at least twice with 8 replications per time.

## Statistical analysis

Means of total and surviving population between treatments were compared using the Glimmix protocol in SAS 9.4 with experimental repeat used as a random factor. Whenever required, post hoc analysis was performed with Tukey’s Honest Significant Difference (Tukey HSD) test at 5% significance level (*P*=0.05). *In planta* enumeration data were analyzed in JMP Pro 14 (SAS Inc.) using Fit Y by X model and One way ANOVA and Tukey HSD at *P*=0.05 after confirming that data were normally distributed and had equal variances.

## Acknowledgements

This research was supported by the following sources for KLH: the USDA National Institute of Food and Federal Appropriations under project PEN04648 and Accession 1015871, the USDA NIFA Foundational Program [2019-67013-29353], and the Lloyd Huck Early Career Professorship.

## Supporting Information

**Fig S1.** Visual representation of the difference in tailocin survival between the stationary and log phase upon 100 AU of tailocin treatment for one hour. St, Stationary phase culture, Log-logarithmic phase culture.

**Fig S2.** Visual representation of the dynamics of persistent and resistant cells after 100 AU of tailocin treatment for 1 and 24 hours. For stationary 1 (St 1), surviving cells did not grow upto 24 hours (indicated by no change in the viable cells 1 and 24 hours post treatment) and were sensitive on re-exposure (i.e. persistent). For stationary 2, although the first hour survivors were sensitive, at 24 hours, population increased due to growth of resistant cells as indicated by their insensitivity to tailocin upon re-exposure (lower panel plates).

**Fig S3.** Testing tailocin activity of supernatants recovered after treatment. Stationary (S) and Log (L) cultures were treated as described above with tailocin for specified number of hours. The treatments were centrifuged, and the supernatant was collected and filter-sterilized. Dilution of the supernatant were tested with *Pph* overlay. Active tailocin particles were recovered from both stationary and log phage treatments at all time points tested (stationary treatments from this experiment gave rise to a lot of tolerant colonies, log had few). Two independent experiments were performed with similar results.

**Fig S4.** Test of tailocin activity after mixing purified tailocin in dilutions of filter-sterilized stationary- and log-phase culture supernatants. Although a slight inhibition of tailocin activity was observed in undiluted stationary supernatants (left), no inhibition was observed after diluting the supernatant (100-1000 fold) as was done for tailocin treatment of cells. After mixing with tailocin, supernatants were incubated for one hour. Dilutions (as shown in the left panel) of tailocin and supernatant mixtures were spotted on a *Pph* overlay. Experiment was repeated twice with three biological replicates and two technical replicates per time (n=12 in total).

**Fig S5.** SDS-Page separation of LPS extracted from wild-type *(Pph),* tailocin high-persistent like mutant of *Pph* (HPL), incomplete resistant (IR) and resistant (R) mutants of *Pph*. IR 7 was one of the incomplete resistant mutants not included in other analyses. Band sizes indicate molecular weight in kilodalton.

**Fig. S6.***In planta* characterization of selected tailocin resistant mutants together with the complemented strains (mutant allele was swapped to WT and vice-versa). Results from two separate experiments with 8 replication per experiment are presented. All complete mutant phenotypes at 24 and 48 hpi had significantly lower population levels compared to WT and incomplete mutant.

**Fig S7.** Growth curve of *Pph* in KB. Cells were inoculated in in a 96-well culture plate with KB and optical density was measured at 600nm every two hours until 24 hourrs. Growth curve experiment was performed twice. One representive experiment is shown.

