## Supplemental material for "*Pseudomonas* can survive bacteriocin-mediated killing via a persistence-like mechanism"

**Supplemental figures**

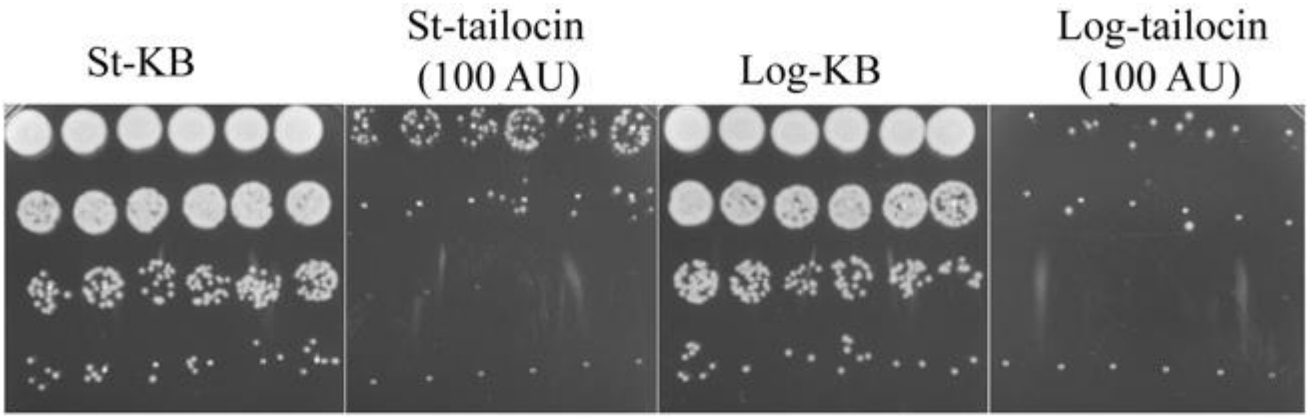


**Fig S1**. Visual representation of the difference in tailocin survival between the stationary and log phase upon 100 AU of tailocin treatment for one hour. St, Stationary phase culture, Log-logarithmic phase culture.

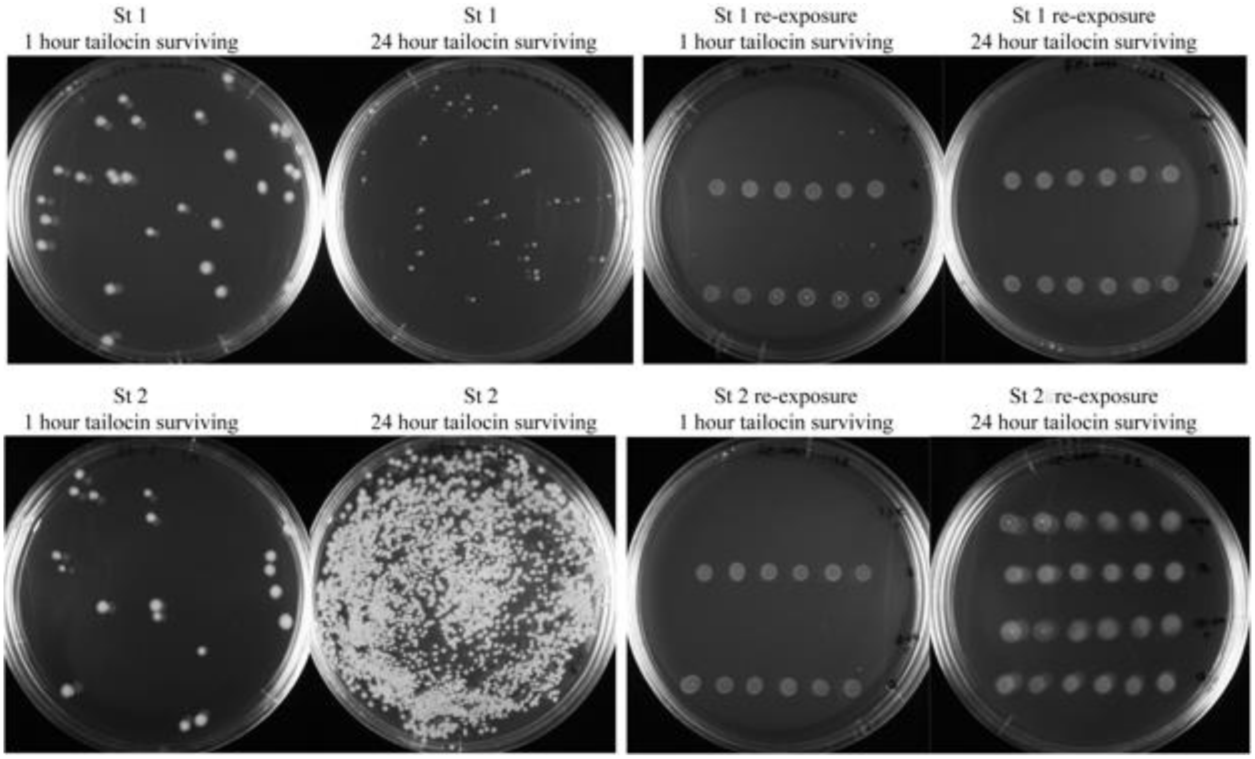


**Fig S2**. Visual representation of the dynamics of persistent and resistant cells after 100 AU of tailocin treatment for 1 and 24 hours. For stationary 1 (St 1), surviving cells did not grow upto 24 hours (indicated by no change in the viable cells 1 and 24 hours post treatment) and were sensitive on re-exposure (i.e. persistent) . For stationary 2, although the first hour survivors were sensitive, at 24 hours, population increased due to growth of resistant cells as indicated by their insensitivity to tailocin upon re-exposure (lower panel plates).

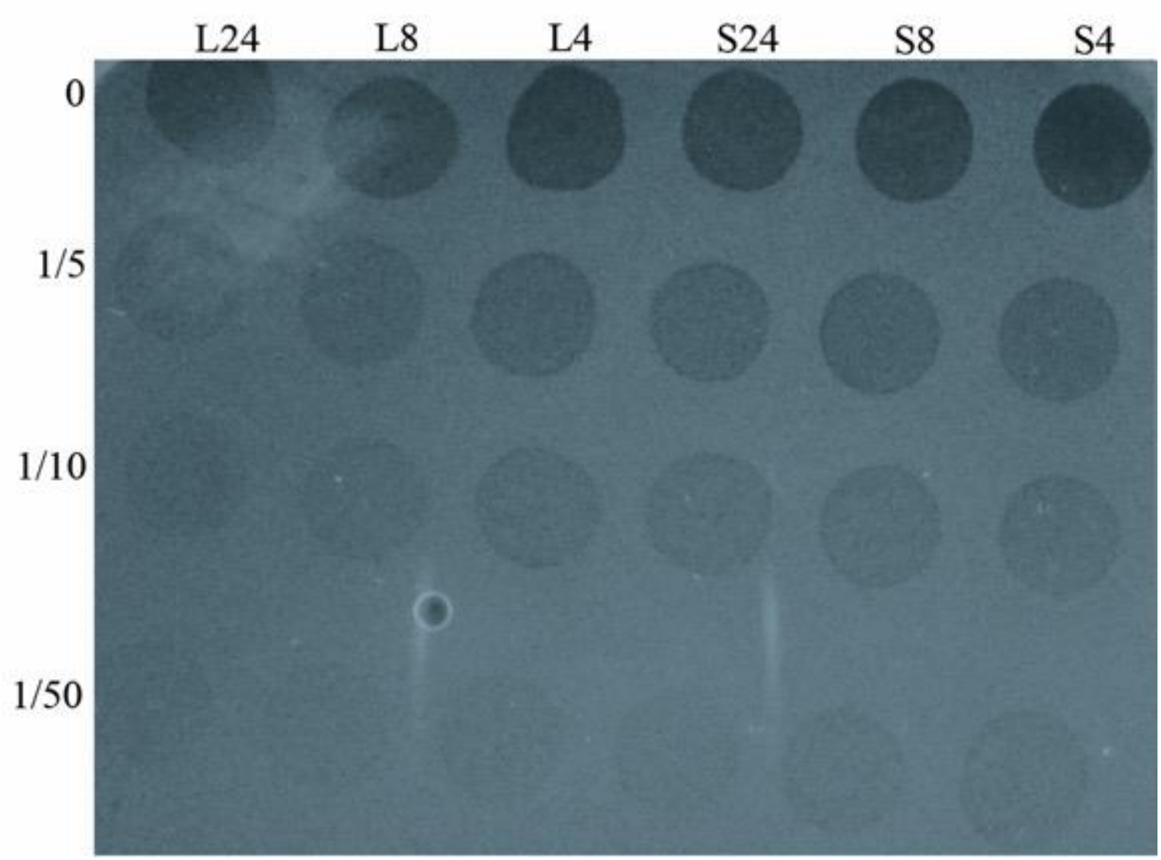


**Fig S3**. Testing tailocin activity of supernatants recovered after treatment. Stationary (S) and Log (L) cultures were treated as described above with tailocin for specified number of hours. The treatments were centrifuged, and the supernatant was collected and filter-sterilized. Dilution of the supernatant were tested with *Pph* overlay. Active tailocin particles were recovered from both stationary and log phage treatments at all time points tested (stationary treatments from this experiment gave rise to a lot of tolerant colonies, log had few). Two independent experiments were performed with similar results.

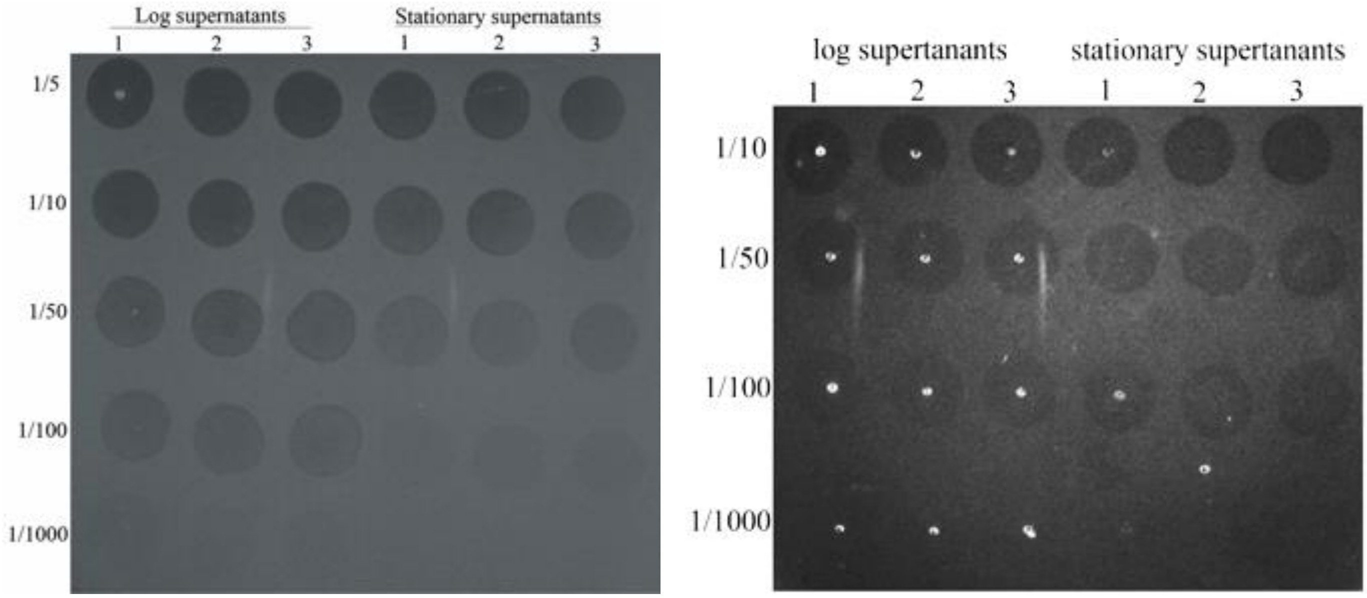


**Fig S4.** Test of tailocin activity after mixing purified tailocin in dilutions of filter-sterilized stationary- and log- phase culture supernatants. Although a slight inhibition of tailocin activity was observed in undiluted stationary supernatants (left), no inhibition was observed after diluting the supernatant (100-1000 fold) as was done for tailocin treatment of cells. After mixing with tailocin, supernatants were incubated for one hour. Dilutions (as shown in the left panel) of tailocin and supernatant mixtures were spotted on a *Pph* overlay. Experiment was repeated twice with three biological replicates and two technical replicates per time (n=12 in total).

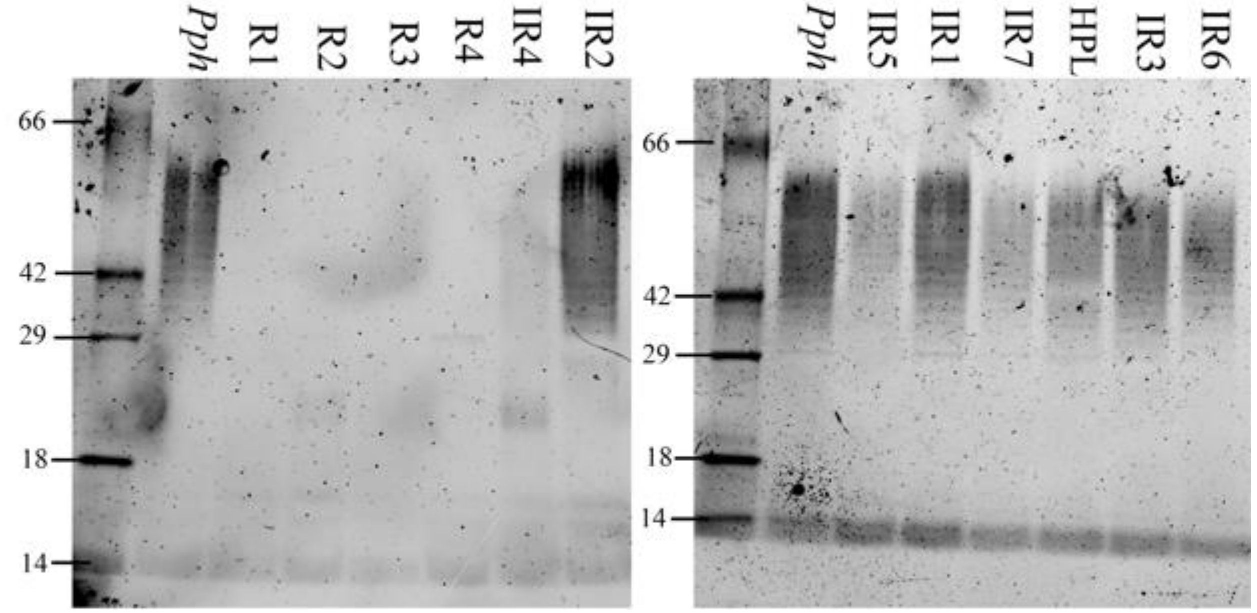


**Fig S5.** SDS-Page separation of LPS extracted from wild-type *(Pph)*, tailocin high-persistent like mutant of *Pph* (HPL), incomplete resistant (IR) and resistant (R) mutants of *Pph*. IR 7 was one of the incomplete resistant mutants not included in other analyses.

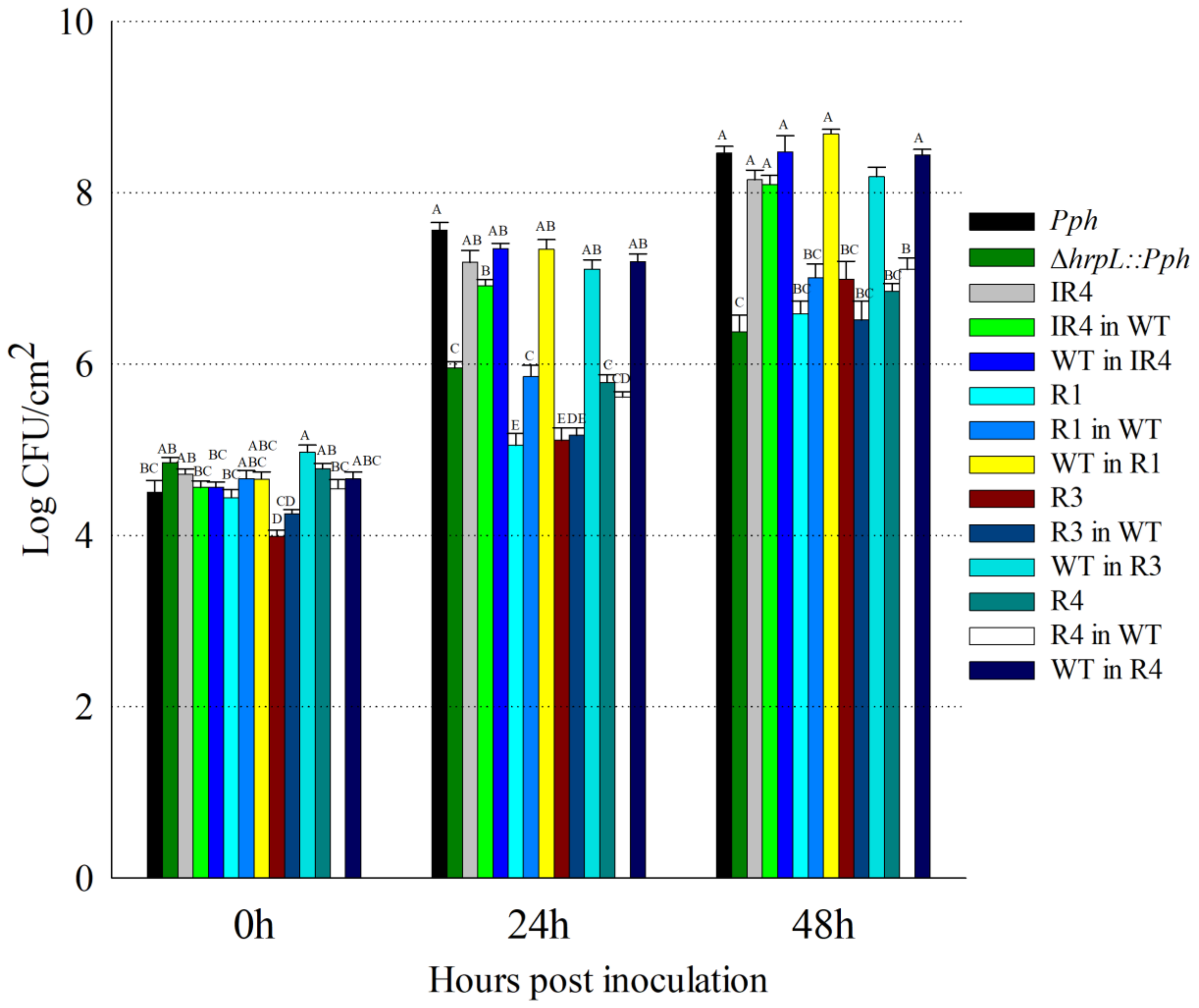


**Fig. S6**. *In planta* characterization of selected tailocin resistant mutants together with the complemented strains (mutant allele was swapped to WT and vice-versa). Results from two separate experiments with 8 replication per experiment are presented. All complete mutant phenotypes at 24 and 48 hpi had significantly lower population levels compared to WT and incomplete mutant.

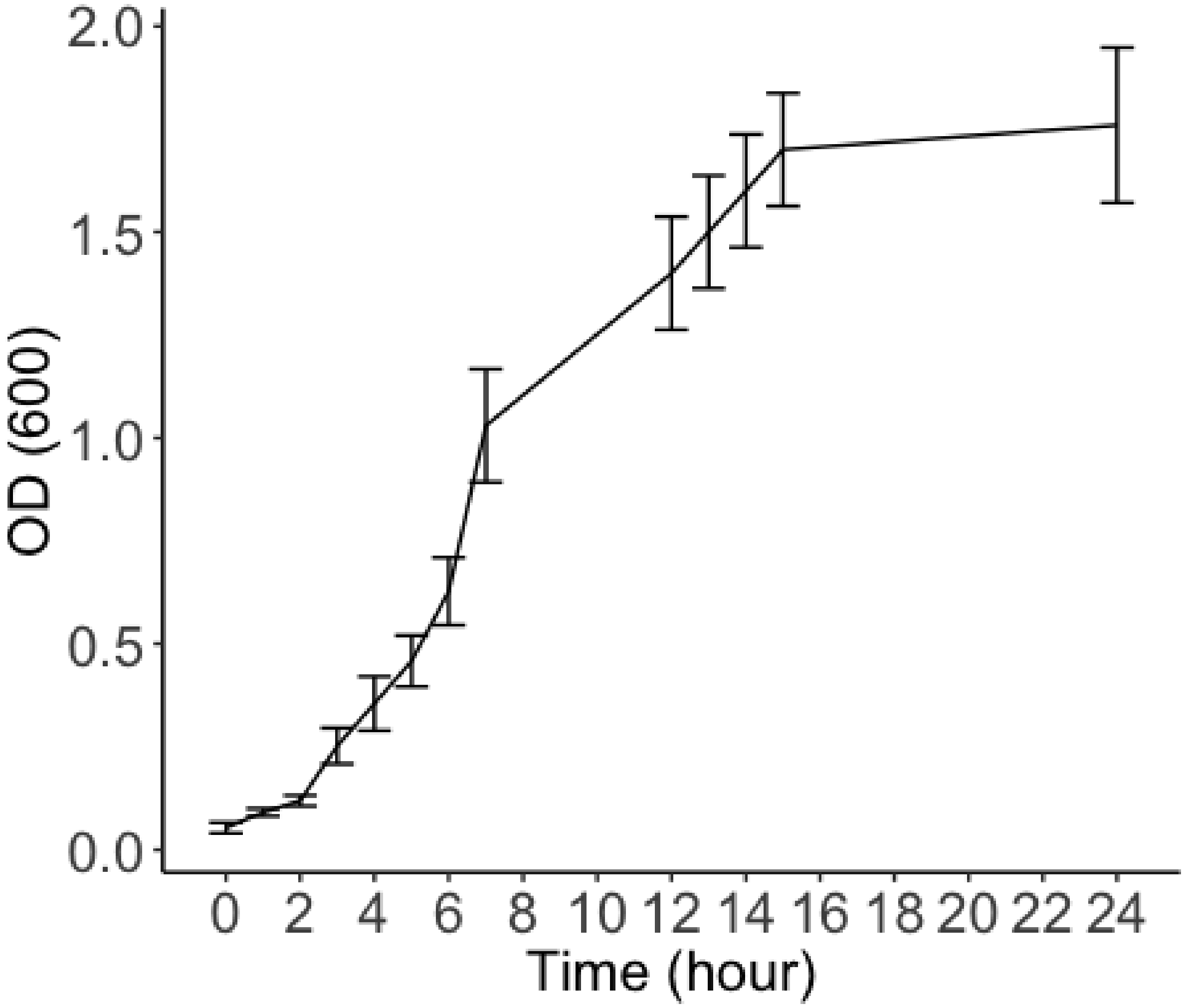


**Fig S7.** Growth curve of *Pph* in KB. Cells were inoculated in in a 96-well culture plate with KB and optical density was measured at 600nm every two hours until 24 hourrs. Growth curve experiment was performed twice. One representive experiment is shown.
